# PCR-based survey of methane-cycling archaea in methane-soaked subsurface sediments of Guaymas Basin, Gulf of California

**DOI:** 10.1101/2023.09.27.559568

**Authors:** John E. Hinkle, Paraskevi Mara, David Beaudoin, Virginia P. Edgcomb, Andreas P. Teske

## Abstract

The Guaymas Basin in the Gulf of California is characterized by active seafloor spreading, rapid deposition of organic-rich sediments, steep geothermal gradients, and abundant methane of mixed thermogenic and microbial origin. Subsurface sediment samples from eight drilling sites with distinct geochemical and thermal profiles were selected for DNA extraction and PCR amplicifation to explore the diversity of methane-cycling archaea in the Guaymas Basin subsurface. We performed PCR amplifications with general (mcrIRD), and ANME-1 specific primers that target the alpha (α) subunit of methyl coenzyme M reductase (*mcrA*). Diverse ANME-1 lineages associated with anaerobic methane oxidation were detected in seven out of the eight drilling sites, preferentially around the methane-sulfate interface, and in several cases showed preferences for specific sampling sites. Phylogenetically, most ANME-1 sequences from the Guaymas Basin subsurface were related to marine mud volcanoes, seep sites, and the shallow marine subsurface. The most frequently recovered methanogenic phylotypes were closely affiliated with the hyperthermophilic *Methanocaldococcaceae*, and found at the hydrothermally influenced Ringvent site. The coolest drilling site, in the northern axial trough of Guaymas Basin, yielded the greatest diversity of methanogen lineages. Our survey indicates potential for extensive microbial methane cycling within subsurface sediments of Guaymas Basin.

## INTRODUCTION

The Guaymas Basin is a young marginal rift basin in the Gulf of California characterized by active seafloor spreading, volcanic sill intrusions into rapidly deposited organic-rich sediments, and steep geothermal gradients where sedimentary organic material of photosynthetic origin turns into hydrocarbons, which can potentially be utilized by microbes as substrates. The dominant hydrocarbon in Guaymas Basin sediments, methane, is of predominantly thermogenic origin in the hot subsurface and gradually changes to microbially produced methane in the upper, cooler sediments [1–3].

Using seafloor samples collected by submersible, hyperthermophilic methanogens (*Methanocaldococcus*, *Methanopyrus*) have been isolated from hydrothermal hot spots in the southern axial trough of Guaymas Basin [4,5]. PCR-based surveys have detected a wide range of methanogenic lineages in surficial Guaymas Basin sediments [6,7], including members of the obligately methylotrophic Methanofastidiosa lineage [8]. Sulfate-dependent anaerobic methane-oxidizing archaea (predominantly ANME-1 and ANME-2) are widespread in surficial sediments of Guaymas Basin [9–12]. Thermophilic ANME-1 lineages were enriched from hydrothermal sediments of Guaymas Basin at temperatures ranging from 50 to 70°C [13–15], and at 70°C from Pescadero Basin in the southern Gulf of California [16, 17].

In contrast to the well-studied surficial sediments of Guaymas Basin, investigations into the diversity and activity of methane-cycling microorganisms in the sedimentary subsurface of Guaymas Basin are just beginning. Enrichments from subsurface sediments obtained during DSDP Expedition 64 demonstrated viable methanogens within the upper 20 m of the sediment column [18]. High-throughput 16S rRNA gene sequencing surveys of the shallow Guaymas Basin subsurface, down to 5 m sediment depth, detected mainly ANME-1 archaea, and found only traces of diverse methanogen lineages [2]. The scarcity of methanogens was also noted by a recent metagenomic study of deep subsurface sediments in Guaymas Basin [3].

To examine the diversity and distribution of methanogens and methane-oxidizing archaea, we performed PCR amplification and sequencing of the *mcrA* gene, which encodes the alpha (α) subunit of methyl coenzyme M reductase, also known as MCR complex. The MCR complex is the central enzyme in anaerobic microbial methane metabolism, as it catalyzes the final step of methanogenesis, and the first step of sulfate-dependent oxidation of methane performed by by methane-oxidizing archaea (anaerobic methane oxidation; 19). While *mcrA* genes are diagnostic for methane-cycling archaea, they also complement the phylogenetic analysis of methane-cycling archaea by 16S rRNA genes [20, 21]. Previous 16S rRNA and *mcrA*-based surveys of surficial Guaymas sediments yielded diverse methanogens and methane-oxidizers, using general and group-specific PCR primers [7]. These primers were constructed at a time when *mcrA*-containing archaea outside of Phylum *Euryarchaeota* were unknown, and thus not targeted [22–25]. Here we use the partial *mcrA* gene sequences to construct phylogenetic trees for a detailed study of methane-cycling archaeal diversity in deep subsurface Guaymas sediments that were sampled during International Ocean Discovery Program (IODP) expedition 385 [26].

## METHODS

### Sampling

Eight drilling sites were visited and sampled by R/V *JOIDES Resolution* during Expedition 385 of the International Ocean Discovery Program to Guaymas Basin (15 September– 15 November 2019). The northern axial trough of Guaymas Basin and its off-axis flanking regions, were targeted to explore the diversity, depth range, and in-situ temperature range of methane-cycling archaea in the Guaymas Basin subsurface (Fig.1). Sediment cores were collected, split into smaller sections on board, and frozen at -80 °C by the shipboard science crew. DNA extraction, and successful *mcrA* amplification occurred at different sites, and sediment depths that extended from 0.8 meters below sea floor (mbsf) down to 142 mbsf (Table 1).

**Figure 1.**
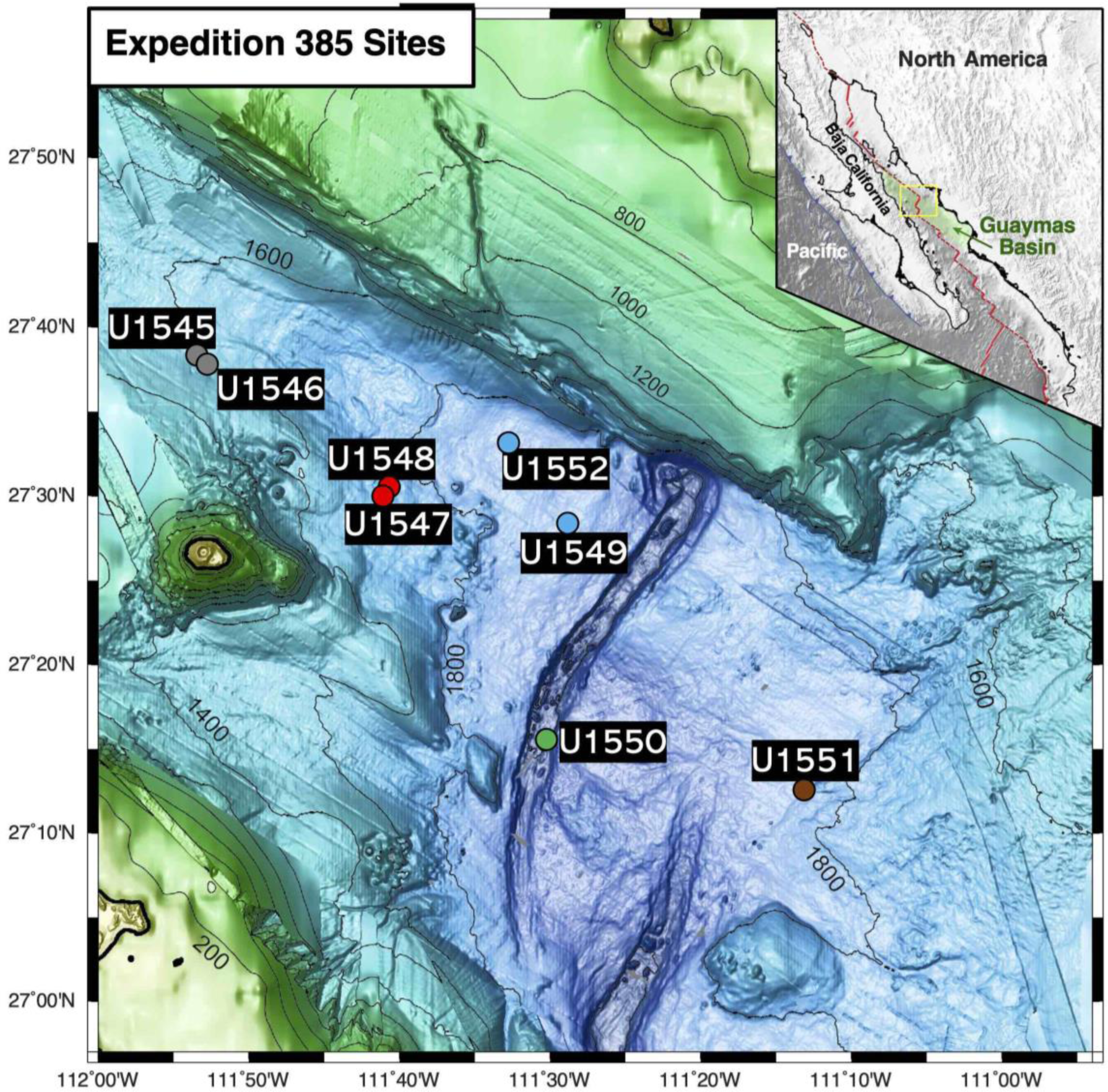
Guaymas Basin drilling site map. The sites follow broadly a northwest-to-southeast transect across the northern Guaymas axial valley. Grey, northwestern off-axis sites U1545 and U1546; red, hydrothermally active Ringvent sites U1547 and U1548; Blue, cold seep sites U1549 and U1552; green, northern Trough axial site U1550; brown, southeastern terrestrially influenced site U1551.

**Table 1.** PCR results for Guaymas Basin subsurface sediment samples. Sample depths are noted as meters below seafloor (mbsf). The (**+**) and (-) symbols indicate successful PCR (+), and no PCR (-) amplifications. A version of this table that includes all samples where PCR amplification was attempted can be found in the supplementary materials (Supplementary Table 1).

| Site | Sample ID | Depth (mbsf) | T (°C) | *mcrA* ANME-1<br>specific primers | *mcrA* general<br>primers (*mcrIRD*) |
| --- | --- | --- | --- | --- | --- |
| U1545B | 1545B_1H2 | 1.7 | 5.3 | — | — |
|  | 1545B_4H3 | 25.8 | 10.7 | + | — |
| U1547B | 1547B_2H2 | 8.6 | 17.4 | — | + |
|  | 1547B_8H2 | 65.7 | 46.6 | + | — |
|  | 1547B_9H2 | 74.2 | 51.0 | + | — |
| U1548B | 1548B_1H2 | 2.1 | 8.2 | + | — |
|  | 1548B_2H3 | 9.1 | 13.7 | + | — |
|  | 1548B_3H4 | 20.4 | 22.9 | + | + |
|  | 1548B_9H3 | 76.5 | 68.0 | + | — |
| U1549B | 1549B_6H3 | 45.6 | 12.1 | + | — |
| U1550B | 1550B_1H2 | 2.0 | 3.8 | + | — |
|  | 1550B_3H2 | 16.9 | 5.8 | + | — |
|  | 1550B_7H2 | 54.8 | 10.9 | + | + |
|  | 1550B_19X2 | 142.0 | 22.7 | + | — |
| U1551B | 1551B_1H1 | 0.8 | 4.8 | + | — |
|  | 1551B_2H2 | 5.8 | 5.3 | + | — |
|  | 1551B_3H2 | 15.4 | 6.3 | + | — |
|  | 1551B_5H2 | 34.2 | 8.2 | + | — |
| U1552B | 1552B_1H2 | 0.8 | 3.8 | + | — |
|  | 1552B_3H3 | 19.2 | 8.6 | + | — |

### Thermal profiles

The Advanced Piston Corer Temperature Tool (APCT-3) [27] and Sediment Temperature Tool (SET-2) were used to measure downhole equilibrium temperatures. The SET-2 tool was used for temperature measurements in lithified sediments that were too stiff for the APCT-3 tool. The SET-2 tool was only used when necessary as it can lead to cracks in the sediment. These cracks lead to temperature measurements of diminished quality. For a detailed protocol of the downhole temperature measurements, refer to Section 3.2 in Neumann et al., 2023. Temperature values were interpolated for each sample using linear regression of the local thermal gradient (°C/m) multiplied by depth (mbsf), plus the y-axis intercept: U1545A,B, T = 0.225 x depth + 4.899; U1546A, T = 0.221 x depth + 2.627; U1547A/B, T = 0.511 x depth + 13.01; U1548A, T = 0.646 x depth + 8.301; U1548B, T = 0.804 x depth + 6.5; U1548C, T = 0.958 x depth + 15.916; U1548D/E, T = 0.271 x depth + 3.241; U1549A, T = 0.194 x depth + 3.228, U1550A, T = 0.135 x depth + 3.532; U1551A, T = 0.1 x depth + 4.741; U1552A/B, T = 0.262 x depth + 3.543.

### DNA extraction

DNA was extracted from near-surface (0.8-2.1 mbsf), intermediate (15.4-54.8 mbsf), and deeper locations (65.7-142 mbsf), as summarized in Table 1. The MP Biomedical FastDNA™ SPIN Kit was used to extract DNA following manufacturer suggestions. Prior to DNA extraction, sediment samples were removed from -80 ^°^C and were thawed on ice. A subsample of 0.5 gr homogenized sediment was transferred using autoclaved and ethanol-rinsed metallic spatula into sterile, DNA/RNA-free tubes containing beads, provided by the manufacturer. All DNA extractions were performed in a UV-sterilized clean hood (two UV cycles of 15 min each) that was installed with HEPA filters. Surfaces inside the hood and pipettes were thoroughly cleaned with RNase AWAY (Thermo Scientific), and with absolute ethanol (200 proof; purity ≥ 99.5%; Thermo Scientific Chemicals) before every extraction and in between extraction steps.

### PCR assays and *mcrA* gene analysis

PCR reactions were performed with *mcrA*-targeting primer sets reported to successfully amplify *mcrA* gene sequences of diverse archaeal methane-cycling lineages identified in Guaymas surficial sediments (7). To PCR-amplify the gene of interest, we used the general mcrIRD primers (F: 5′-TWYGACCARATMTGGYT-3′; R: 5′-ACRTTCATBGCRTARTT-3′) that cover a range of known *mcrA* gene sequences in Guaymas Basin, and the ANME-1-specific *mcrA* gene primers (F: 5′-GACCAGTTGTGGTTCGGAAC-3′; R: 5′-ATCTCGAATGGCATTCCCTC-3′) that target the ANME-1 lineage involved in anaerobic methane oxidation in Guaymas Basin (7). The PCR reactions at every examined depth were performed in triplicate (n=3) using Takara SpeedSTAR HS DNA polymerase kit (Takara Bio USA, Madison, WI) with the following modifications: each 25 µl PCR reaction contained 0.5-1 ng of template DNA, 2× Takara SpeedSTAR Buffer I, 2.5 mM dNTPs (Takara Bio USA, Madison, WI), 2 mM of each primer (final concentrations), 1.25 units of SpeedSTAR HS DNA Polymerase and DEPC water (Fisher BioReagents). The PCR reactions were performed at 95°C for 5 min, followed by 35 cycles of 95°C (30 s), 48°C (30 s), and 72°C (45 s) for the mcrIRD primers. Same PCR conditions were also applied for ANME-1-specific *mcrA* primers with the exception that the annealing temperature was set at 60°C (30 s). To confirm absence of contamination due to handling and PCR reagents, we included negative controls (blanks) in all PCR experiments. Negative controls did not include template DNA, but 5 µl of DEPC water. As positive control we used DNA from surficial Guaymas sediments collected with *Alvin* push cores during the AT42-05 research cruise in Guaymas Basin, containing natural assemblages of methanogens and ANME-1 archaea [28].

2 μl of all PCR amplified samples were run on 2% agarose gel (Low-EEO/Multi-Purpose/Molecular Biology Grade Fisher BioReagents) to verify the amplification of the *mcrA* genes. The PCR-amplified *mcrA* gene using mcrIRD or ANME-1 primers is ∼ 500 base pairs in size. All successful PCR reactions per sample (n=3) were pooled together, and were purified and concentrated using the Agencourt AMPure XP protocol (Beckman Coulter) following the manufacturer’s suggestion. The purified/concentrated PCR products were sequenced at Georgia Genomics and Bioinformatics Core (University of Georgia) using the MiSeq Illumina Platform.

The paired reads generated with MiSeq were quality checked and trimmed using FastQC (v. 0.11.7). We analyzed the *mcrA* data using the QIIME2 platform [29], and the DADA2 plugin provided in the QIIME2 pipeline to denoise and optimize the merging of the forward and reverse reads. The functional annotation of the generated *mcrA* sequences was assigned manually using BLASTx against GenBank NR. The *mcrA* and ANME-1 *mcrA* sequences that were retained for further analyses, were those that were functionally annotated as “methyl CoM reductase” and presented *e* values ≥ 1e-10. Those manually curated sequences that were functionally misannotated were excluded from downstream analyses.

### *mcrA* gene phylogeny

Those *mcrA* gene sequences successfully annotated “methyl CoM reductase” were aligned using Multiple Sequence Comparison by Log-Expectation (MUSCLE) in MEGA11. The aligned sequences were used to construct minimum evolution phylogenetic trees checked by 1000 bootstrap replicates. Bootstrap values indicating greater than 50% support for an individual cluster were included in each phylogenetic tree. For easy reference, all *mcrA* gene sequences in our phylogenetic trees are included in the supplementary materials.

Nucleotide BLAST search (BLASTn) was used to find closely related sequences in GenBank NR. These closely related sequences were then included in the trees to add phylogenetic context and allow for cluster labeling. A phylogenetic tree was constructed using the five most abundant ANME-1 *mcrA* gene sequences from seven of the eight drilling sites. *mcrA* gene sequences from methanogenic lineages were placed into a separate phylogeny. To obtain a wider view of ANME-1 diversity, site-specific trees were constructed for sites where the number of sequences recovered was conducive to constructing a phylogenetic tree. For each site, all sequences with >100 clones were used to construct a phylogenetic tree. An exception was for the site U1550B specific tree which had 200 sequences with >100 clones. For this site, the top 25 sequences, by clone number, from samples U1550B_ 1H2, U1550B_3H2, and U1550B_7H2, (2.0, 16.9, and 54.8 mbsf, respectively; Table 1) along with two sequences from sample U1550B_ 19X2 (142 mbsf) were used to construct a phylogenetic tree.

## RESULTS

### Site characteristics

PCR-amplified *mcrA* genes were obtained from nearly all drilling sites of IODP Expedition 385 (Fig. 1) which differ in their degree of hydrothermally driven heat flow [30], and in the relative extent of marine vs. terrestrial sedimentation [31]. Samples yielding PCR amplicons were generally affiliated with subsurface maxima or with broad zones of high alkalinity and DIC concentration [32], a proxy for microbial organic matter remineralization (Supplementary Figures S1a-h and S2). These geochemical layers often coincide with methane-sulfate transition zones, which are shown to have elevated microbial cell counts and increased microbial activity [33]. Sediment samples, DNA yields, and PCR results are summarized in Table 1. Two neighboring sites (U1545 and U1546) on the northwestern end of Guaymas Basin [34, 35] essentially differ by the presence of a massive, thermally equilibrated sill between 350 to 430 meters below seafloor (mbsf) at Site U1546 [36]. Two drilling sites (U1547, U1548) at the hydrothermally active Ringvent area, approximately 28 km northwest of the spreading center [2] target a shallow, recently emplaced, hot sill that creates steep thermal gradients and drives hydrothermal circulation [37]. Site U1549 [38] samples the periphery of an off-axis methane cold seep, Octopus Mound, located ∼9.5 km northwest of the northern axial graben. Site U1550 is located close to DSDP Site 481 and samples the heterogeneous (marine and terrestrial) sediments that are accumulating in the northern axial valley [39]. Site U1551 represents the terrestrially influenced sediments of the southeastern Guaymas Basin with riverine sand input from the Yaqui River [40]. Site U1552 [41] sampled the subsurface of a cold seep site surrounded by shallow methane hydrates, close to the Sonora Margin.

Regarding their in-situ temperature gradients, the Guaymas Basin drilling sites fall into two categories, the hot Ringvent sites with steep temperature gradients of ca. 50 to 100°C/100 m depth, and the majority of Guaymas sites with elevated temperature gradients of approx. 20 to 25°C/100 m (Fig. 2). Even the gradient at the coolest site, U1550 with ca. 13-14°C/100m, is at least twice as steep as the temperature gradients commonly found for ocean crust. These distinct thermal gradients are to some extent reflected in different lineages of detected methanogens.

**Figure 2.**
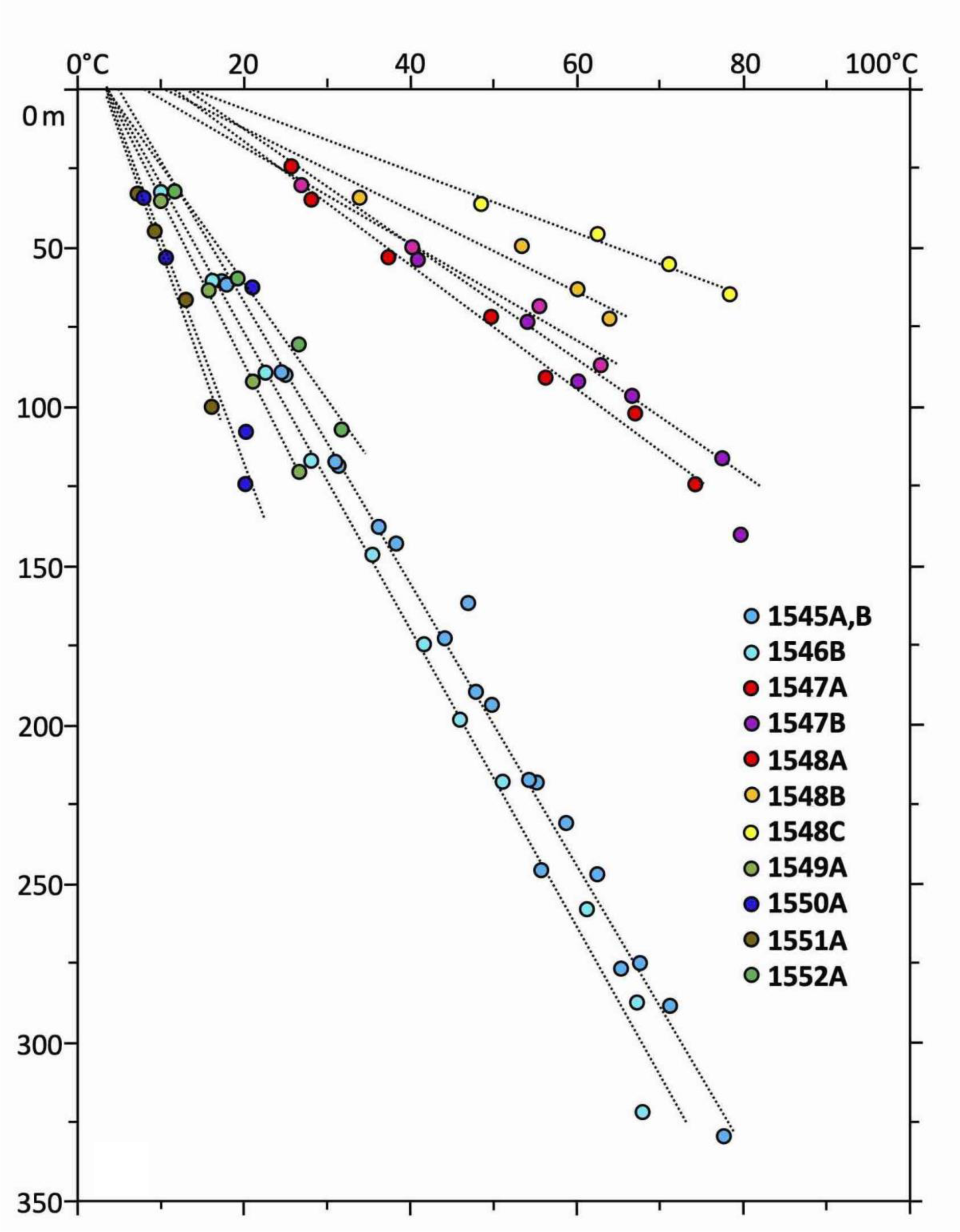
Temperature profiles of Guaymas drilling sites, plotted from measurements tabulated in the site chapters of Expedition 385 [35–41].

### Detection of Methanogen lineages

Three of the seven examined sites (U1547B, U1548B, and U1550B) yielded *mcrA* genes related to previously established methanogens or methane-cycling archaea, using PCR primers that encompass the widest possible diversity of *mcrA* genes, but do not capture ANME-1 [7]. Most of our positive PCR amplicon results were obtained from the upper layer of the sediment column, whereas deeper sediments yielded negative results, even after many PCR attempts (Table 1).

Two sites (U1547B and U1548B) are located at the Ringvent mound, an off-axis hydrothermal system driven by recent sill emplacement (Fig. 3). Here, the temperature gradients increase rapidly along a sampling transect with tightly spaced, increasing temperature measurements towards the ring-shaped mound (Fig. 2) where conspicuous hydrothermal activity and a topographical high of ca. 20 m mark the rim of the buried sill [2]. At Ringvent sites U1547B and U1548B, the most abundant phylotypes were close relatives of the extreme hyperthermophile and obligate hydrogenotroph *Methanocaldococcus bathoardescens* (Fig. 4), originally isolated from low-temperature hydrothermal fluid (26°C) at the Axial Seamount on the Juan de Fuca Ridge [42]. Although isolated from relatively temperate fluids, *Methanocaldococcus bathoardescens* is a hyperthermophile that grows under temperatures from 48 to 90°C, with a thermal optimum of 82°C [42]. Samples from the Ringvent mound sites also yielded *mcrA* genes of the *Methanosarcinaceae* (capable of using acetate, CO_2,_ and methyl substrates for methanogenesis) and the mostly hydrogenotrophic *Methanomicrobiaceae* (Fig. 4).

**Figure 3.**
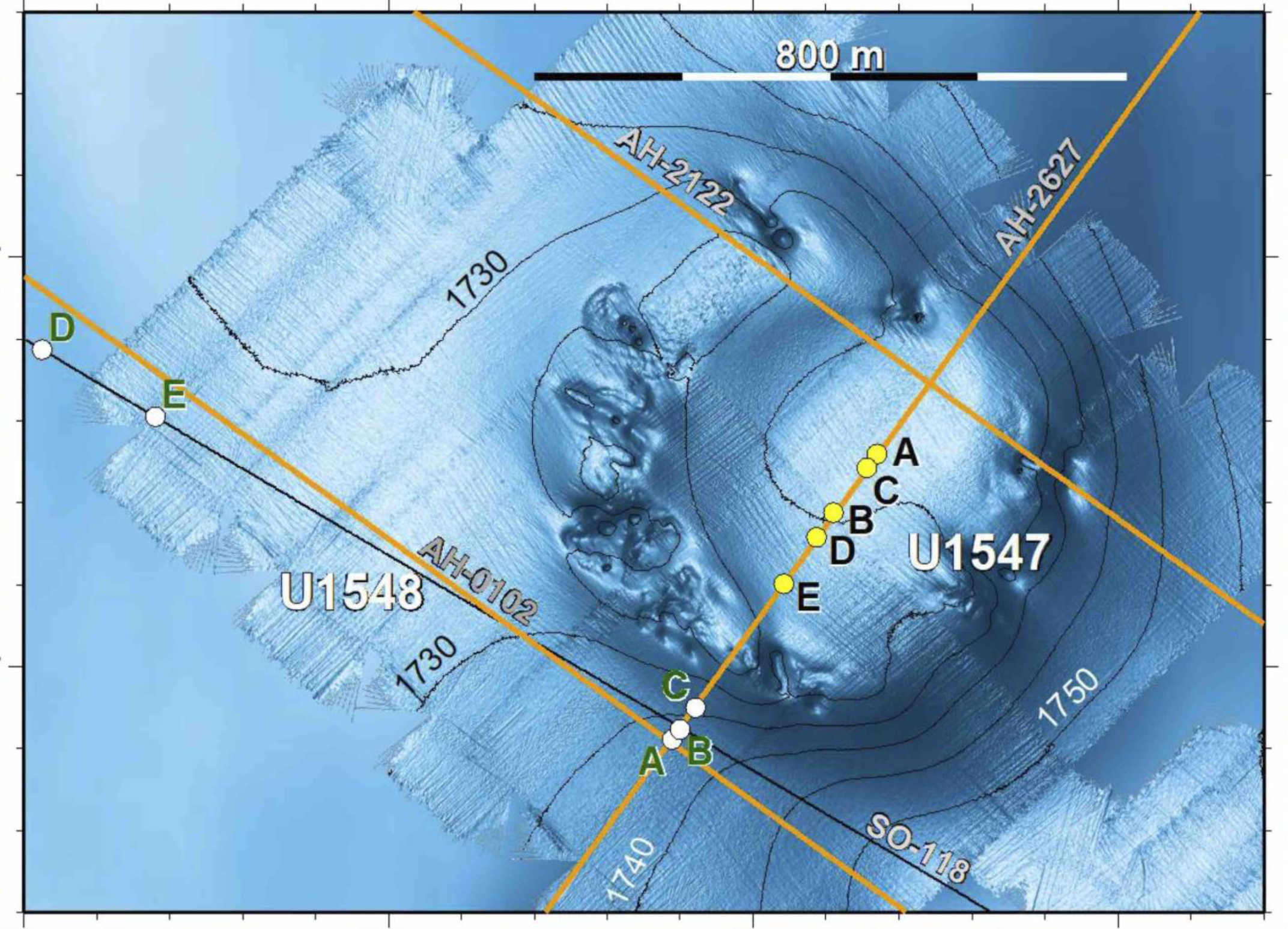
Ringvent Mound site with IODP Expedition 385 drilling sites and hole positions (U1547A-E and U1548A-E), and multichannel seismic lines for subsurface mapping [37] The topographical high marks the edge of the shallow subsurface sill where hydrothermally circulating fluids reach the sediment surface. Sequence data from cores U1547B and U1548B represent the microbial communities within the interior sedimented bowl, and the hydrothermally impacted communities just outside the sill.

**Figure 4.**
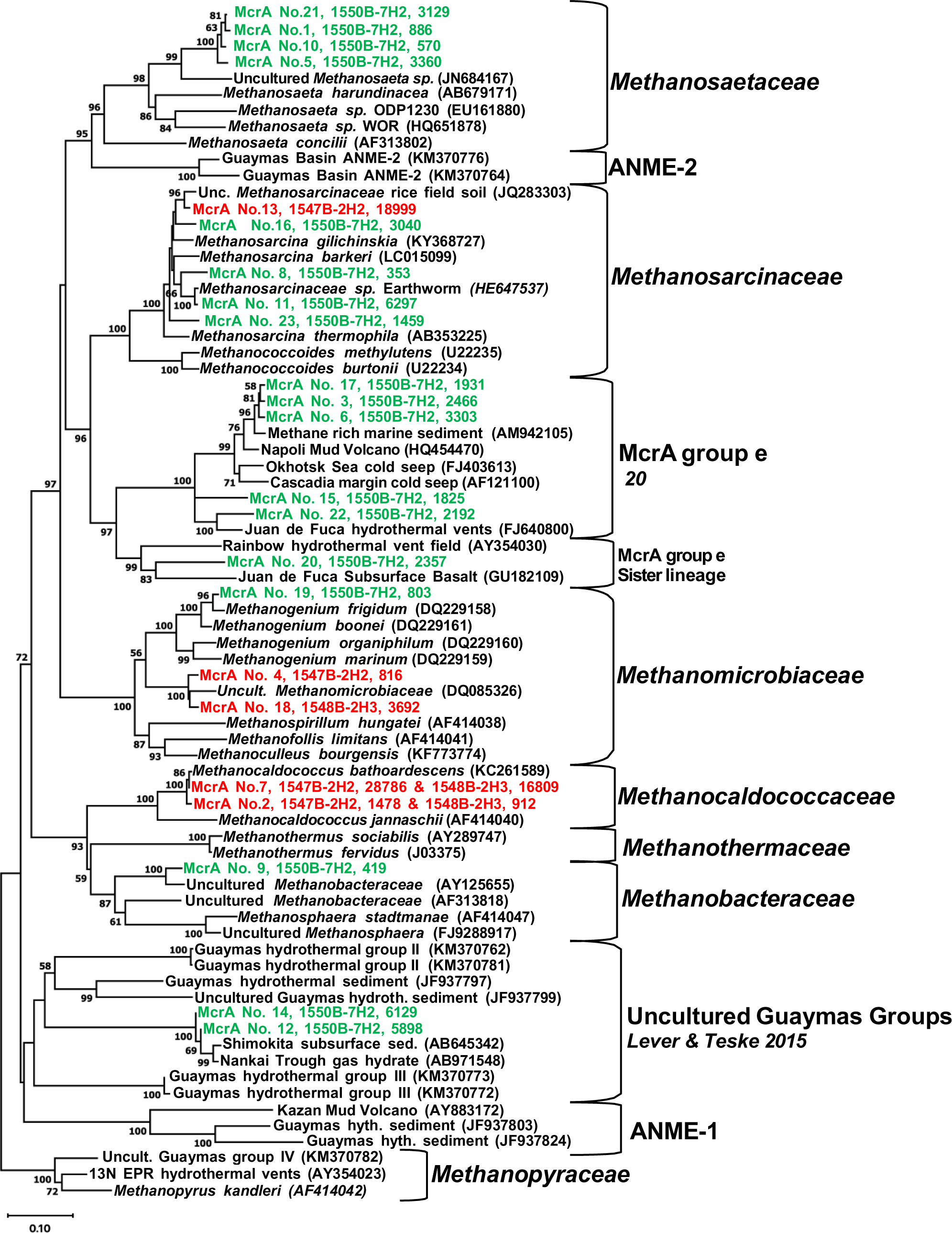
Phylogenetic diversity of methanogen lineages detected with general methanogen primers, resulting in 23 ASVs of partial *mcrA* gene sequences. The phylogeny was constructed using the minimum evolution method of phylogenetic inference and included bootstrap values > 50%. ASVs from Ringvent sites U1547B and U1548B are plotted in red, and ASVs from axial site U1550B are plotted in green. Each taxon label starts with the ASV number, followed by the IODP sample code (drilling site, subcore number and segment), and the number of sequences within each ASV. *Methanocaldococcaceae* ASVs No. 2 and 7 were recovered from both samples each, U1547B_2H2 and U1548B_2H2.

At the cool axial site, U1550B, greater methanogenic diversity was observed, including the obligately acetoclastic *Methanosaetaceae*, the metabolically diversified *Methanosarcinaceae*, and the most hydrogenotrophic or (occasionally) ethanol-utilizing *Methanomicrobiaceae* and *Methanobacteraceae* (Fig. 4). Additionally, the uncultured *mcrA* “group E” lineage related to anaerobic methane-oxidizing ANME archaea [21], and an uncultured Guaymas group without any cultured relatives were detected at this site [7].

### ANME-1 diversity

While methanogen populations were only observed at three sites (U1547B, U1548B, and U1550B), seven sites yielded *mcrA* gene sequences affiliated with the ANME-1 lineage (Methanophagales), using primers fine-tuned for this order-level lineage (7). Extensive searches in GenBank NR and comparisons to database sequences showed that ANME-1 phylotypes from Guaymas Basin subsurface samples were closely related to reference sequences from the Napoli mud volcano [43], from subsurface sediments of the Mississippi Canyon 118 seep site in the Gulf of Mexico [44], and from shallow subsurface sediments recovered by piston coring and push coring in Guaymas Basin [2, 45]. Almost all ANME-1 *mcrA* gene sequences could be accommodated in sixteen distinct phylogenetic clusters (Figure 5). These clusters were further examined and confirmed through site-specific *mcrA* gene phylogenies for five sites, U1547B, U1548B, U1550B, U1551B, and U1552B (Supplementary Figures S3 to S7).

**Figure 5.**
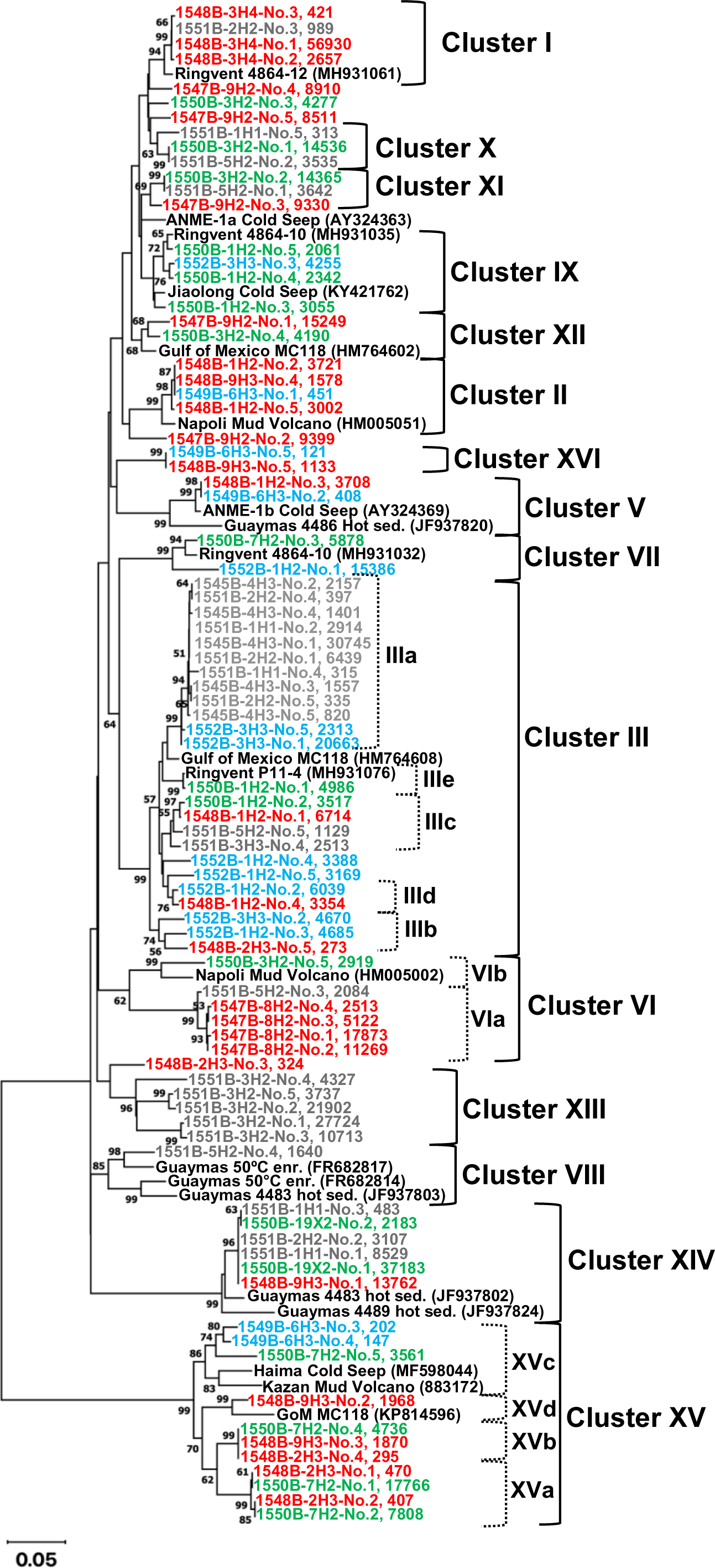
General ANME-1 minimum evolution phylogeny based on partial *mcrA* gene sequences, encompassing the five most frequently recovered ASVs from each drilling site and sample. Reference site U1545B sequences are plotted in grey, those from Ringvent sites U1547B and U1548B are plotted in red, axial site U1550B sequences appear in green, terrestrial site U1551B sequences in brown, and seep site U1549B and U1552B sequences are plotted in blue. Each taxon label starts with the IODP sample code (drilling site, subcore number and segment), followed by the ASV number and the number of sequences within each ASV. Site-specific phylogenies that include a greater number of ASVs (more than five) for each site are available as Supplemental Figures S2 to S6.

In searching for thermophilic ANME-1 lineages, we found that cluster VIII, consisting of *mcrA* genes from ANME-1 enrichments in surficial sediments at 50°C [13], also included Guaymas subsurface *mcrA* genes from site U1551B. However, we observed no overlap of Guaymas subsurface *mcrA* genes with *mcrA* genes from thermophilic methane-oxidizing ANME-1 enrichment cultures that are active at 70°C [15, 17].

Habitat and site preferences of different *mcrA* gene lineages were visualized with a bubble plot of *mcrA* gene recovery, color-coded by habitat type (Fig. 6, and Supplementary Figure S8). From the relatively cool northwestern site: U1545B, only a single sample yielded *mcrA* gene sequences, all of them members of cluster 3 (color-coded in grey). Gene sequences from the hydrothermal Ringvent sites (in red, Fig. 6) were highly diverse, and affiliated with clusters 1, 2, 5, 6, 10, 11, 12, 14, 15 and 16. Gene sequences from the axial site U1550B were also diverse (color-coded in green, Fig. 6). Gene sequences from the terrestrially influenced site U1551B were largely affiliated with group 13 (Fig. 5), plus smaller proportions of U1551B-derived sequences that fell into clusters 1, 3, 8 and 10 (Fig. 6). Cluster 13, which dominated the samples from site U1551, did not include any sequences from other sites and thus provides the best example, within this dataset, for strong habitat preference within a particular lineage (color-coded in brown, Fig. 6). No published *mcrA* genes that would fall into this cluster were found in GenBank NR. Finally, gene sequences from the cold seep site U1552B were affiliated with clusters 3, 7, and 9 (color-coded in blue, Fig. 6).

**Figure 6.**
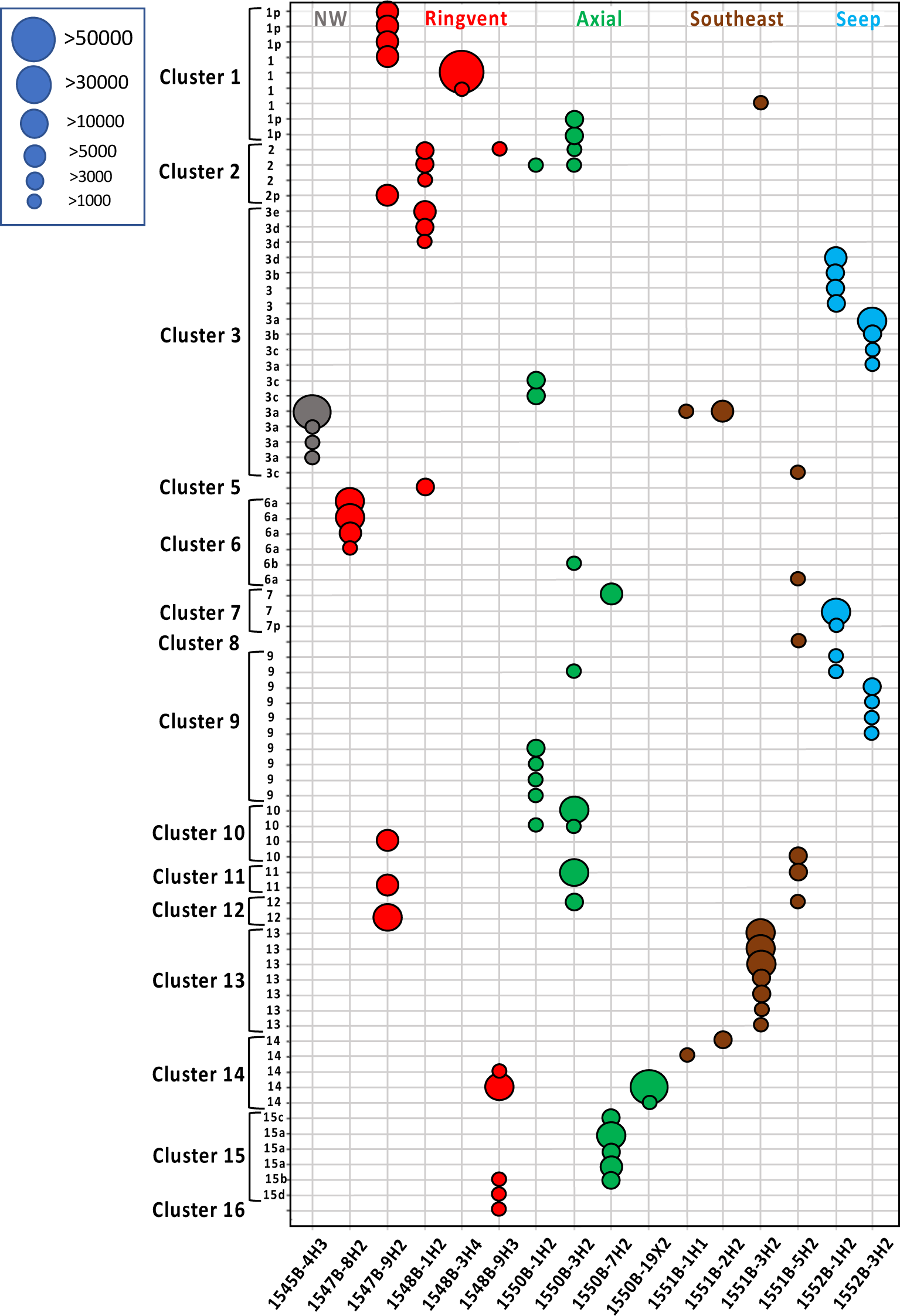
Abundance dot plot of mcrA gene amplicons (represented by > 1000 sequences) color-coded by site, as in Figures 1, 4 and 5. Dot size indicates number of sequences. The x-axis indicates the samples, and the y-axis indicates the *mcrA* sequences and their clustering. A version with full sample annotation on the left margin is available as Supplementary Figure S8. ASVs that are only peripherally associated with specific mcrA gene lineages, lacking bootstrap support, are labelled with “p”.

## DISCUSSION

### ANME-1 diversity

The methane-oxidizing group ANME-1, is widespread in the Guaymas Basin, as it was observed at every site examined, particularly near depths where methane-sulfate interfaces occur. Methane-sulfate interfaces are present at varying depths in the subsurface, depending on geologic formations and methane flux. To some extent, the different regions in the Guaymas Basin are characterized by different ANME-1 types. While it is possible that some clades may not be detected due to sensitivity issues or sequencing artifacts, our data identify clades that show an observable preference for particular sites, which differ in temperature and geochemistry. This differentiation can be observed by comparing the clusters in the phylogenies constructed specifically for each drilling site. For example, when comparing the ANME-1 phylogenies from sites U1548B and U1552B, the methane hydrate site U1552B contains three distinct ANME-1 clusters, two of which are not shared with the phylogeny of hydrothermally influenced Ringvent site U1548B (Figs. S4 & S8 in the supplemental material). These differences in ANME-1 cluster representation across sites suggest how the locally distinct thermal profiles (Fig. 2) and in-situ temperature conditions contribute to shaping the microbial biogeography of the Guaymas Basin.

### Mud volcano connection

Close phylogenetic relationships between sequences from mud volcano and deep subsurface sediments were observed in the general ANME-1 phylogeny (Fig. 5). Previous research indicates that these similarities reflect the role of mud volcanoes as a “window to the deep subsurface”, dispersing microbes from the deep subsurface into the hydrosphere; as mud volcanoes erupt they bring gas-rich fluidized sediment and its microbial populations from the deep subsurface, to the seafloor [46]. Mud volcanoes could garner more attention as an alternative way to study the deep marine biosphere since R/V *JOIDES Resolution* will be decommissioned in October 2024 and plans for a replacement vessel have been paused for a prolonged period [47].

### Hyperthermophilic methanogens at Ringvent sites

Hyperthermophilic methanogens (family *Methanocaldococcaceae*) at the hot Ringvent sites U1547B (51°C at 74.2 mbsf) and U1548B (68°C at 76.5 mbsf), were not recovered from deep, hot sediments, but were recovered from shallow, cool, and relatively oxidizing (sulfate-reducing) sediment samples between 2.1 and 9.1 mbsf (up to 17.4°C). This apparent paradox can be explained by the low cell abudnancies (≤ 10^5^ cells per cm^3^ of sediment) detected at those two sites when sediments reached temperatures ≥ 45°C [48], and by the environmental regime that exists in the Ringvent area: The Ringvent mound is underlaid by a hot volcanic sill, which drives hydrothermal circulation at the relatively thinly sedimented surface of the mound, and brings hot hydrothermal fluids (up to 75 °C) to the sediment surface [2]. Hyperthermophiles could rise with methane-rich hydrothermal fluids from deeper sediments, or may originate from localized hydrothermal hot spots that have been observed at the sediment/water interface of Ringvent [2]. Either way, hydrothermal circulation could deposit microbial cells in the wider Ringvent area and seed cool, surficial sediments with hyperthermophiles. Previous studies have detected thermophilic methanogen lineages in cool, surficial Guaymas Basin sediments, and have proposed transport to the surface via hydrothermal flow [6].

### Methanogen scarcity

Many ANME-1 sequences were collected from our sampling sites; however, methanogen sequences were relatively scarce. This poses a question: Why are ANME-1 so abundant, yet methanogens so rare in the Guaymas Basin deep subsurface?

While methodological limitations, such as PCR primer range and sensitivity, could play a role, methanogen scarcity is not only observed in this *mcrA* gene survey, but also in 16S rRNA analyses [2, 12], in metagenomic investigations [3], and in cultivation studies [49]. It seems unlikely that these conceptually and technically distinct methods are consistently biased against methanogens, and to such a degree that they would miss a large swath of methanogenic diversity. While undetected, “exotic” methanogenic lineages cannot be excluded with certainty, they would have to hide in plain sight (for example, in poorly-described uncultured phylogenetic lineages) to comprise a significant portion of methanogenic diversity.

The likely scarcity of methanogens in the Guaymas Basin deep subsurface has implications for the origin and microbial processing of subsurface methane. Methane clumped isotope measurements show that the proportion of biogenic methane increases towards the sediment surface, whereas methane in deep, hot sediments is predominantly thermogenic [3]. If the population density of active methanogens in the Guaymas subsurface sediments is relatively low, the massive reservoir of biogenic subsurface methane must have accumulated over geological time [3]. Methanogenic gene expression is predominately observed in surficial sediments where most methane-cycling occurs [50]. Some of this methane produced in surficial sediments could be incorporated into the subsurface reservoir of biogenic methane.

### Thermal preferences of methanogens

Notably, the cooler axial site (U1550B) exhibits greater methanogenic diversity than the much warmer Ringvent mound sites (U1547B and U1548B). This difference in diversity may reflect the methanogens’ general preference for moderate temperatures [12]. While thermophilic methanogenic lineages have been isolated from deep-sea vent systems [4, 42], most cultured methanogenic lineages are not thermophilic [51]. Observing greater methanogenic diversity in cooler sediments of the Guaymas subsurface underscores the linkage between sediment depth/temperature and methane origin, by supporting the observation that biogenic methane predominates in shallower, cooler sediments, whereas thermogenic/abiotic methane predominates in deeper, warmer sediments [3].

### Future directions

A future direction of this research lies in investigating the matching 16S rRNA datasets and pulling *mcrA* genes from entire genomes. *mcrA* genes pulled from previously published genomes will allow for the strengthening of our current *mcrA* phylogenies. The construction of 16S rRNA phylogenies to complement the existing *mcrA* phylogenies will allow for a comprehensive PCR-based survey of methane-cycling microbial communities in the Guaymas Basin subsurface. Additionally, there is the possibility of linking our *mcrA*-based phylogenies to genome-based phylogenies of ANME-1 [17].

The diversity of Guaymas Basin subsurface sediments suggests that they would contain *mcrA*-containing archea outside of Phylum *Euryarchaeota*. Candidates could even be hiding in published metagenomes, for example, metagenomes related to Phylum *Verstraetearchaeota* [52]. Further development of *mcrA* primers to capture these groups would aid in their detection [23, 53].

Another research direction focuses on the volcanic sills of Guaymas Basin. Current research suggests that the abundance of methane located at and in these sills is of predominately thermogenic origin, resulting from the thermal maturation of organic matter [1,3,37]. However, these sills are known to harbor sediment intrusions, which could contain microbes, and even methanogens. The sediment-sill interfaces remain within the temperature range for methanogenic hyperthermophiles [5]. While it is unknown if there is enough accessible energy to support cellular metabolism, sill intrusion contributes to the thermal maturation of organic matter and provides a potential influx of low-molecular weight maturation products that would be available for microbial metabolism [54, 55]. Future metagenomic analyses of recovered veined rock samples will seek to address this issue.

## DATA AVAILABILITY

All *mcrA* generated sequences were deposited into the National Center for Biotechnology Information (NCBI) Sequence Read Archive database under project BioProject ID PRJNA909197. The accession numbers for the *mcrA* sequences from samples U1550B_7H2, U1548B_2H3, and U1547B_2H2 are SRR23604134, SRR23604145, and SRR23604146 respectively. The accession number for the ANME-1 *mcrA* sequences from sample U1545B_4H3 is SRR23604132. The accession number for the ANME-1 *mcrA* sequences from samples 1547B_8H2 and 1547B_9H2 are SRR23604131 and SRR23604130, respectively. The accession numbers for ANME-1 *mcrA* sequences from samples 1548B_1H2, 1548B_2H3, 1548B_3H4 and 1548B_9H3 are SRR23604128, SRR23604129, SRR23604127 and SRR23604126, respectively. The accession number for the ANME-1 *mcrA* sequences from sample 1549B_6H3 is SRR23604144. The accession numbers for ANME-1 *mcrA* sequences from samples 1550B_1H2, 1550B_3H2, 1550B_7H2 and 1550B_19X3 are SRR23604143, SRR23604142, SRR23604141 and SRR23604140, respectively. The accession numbers for ANME-1 mcrA sequences from samples 1551B_1H1, 1551B_2H2, 1551B_3H2 and 1551B_5H2 are SRR23604139, SRR23604138, SRR23604137 and SRR23604136, respectively. The accession numbers for ANME-1 mcrA sequences from samples 1552B_1H2 and 1552B_3H3 are SRR23604135 and SRR23604133, respectively.

## AUTHOR CONTRIBUTIONS

JEH wrote the first draft of the manuscript, conducted the *mcrA* phylogenetic analyses, and created phylogeny figures. PM conducted the *mcrA* PCR amplifications, and the manual curation and functional annotation of the *mcrA* sequence data. DB conducted DNA extractions and the *mcrA* sequence data analyses. VE collected sediment samples, and supervised PM and DB on the *mcrA* data generation and analyses. AT conceptualized the study design, provided figures, and contributed to subsequent drafts of the manuscript. All authors contributed to and approved the final form of this draft.

## CONFLICT OF INTEREST

Authors declare no conflict of interest.

## ACKNOWLEDGMENTS

Sampling in Guaymas Basin and post-cruise research in the Teske lab was supported by the International Ocean Discovery Program. We thank David Geller-McGrath for an early version of Figure 6. This study was supported by NSF Grant OCE-2046799 to VE, PM, AT, and R. Hatzenpichler, and by NSF grant OCE-1829903 to VE, PM, and AT. Current research in the Teske lab is supported by NASA Exobiology grant 512580 and by NSF Biological Oceanography grant 5120047.

## Notes

### Competing Interest Statement

The authors have declared no competing interest.

### Summary of Updates

Phylogenetic trees revised (thicker branches, larger text on tables). Supplemental Figure 1 adjusted to increase readability. Minor additions to text of manuscript to reference mcrA-containing archaea outside of Phylum Euryarchaeota.

