## Supplemental Materials for "PCR-based survey of methane-cycling archaea in methane-soaked subsurface sediments of Guaymas Basin, Gulf of California"

**Supplementary Table 1**. PCR results for Guaymas Basin subsurface sediments using general *mcrA* primers (mcrIRD; this study), *mcrA* specific primers (ANME-1; this study), and primer combination of 25F and 806R for archaeal 16S rRNA (Mara et al., 2023). Sample depths are noted as CSF-A depths (IODP 2011). A “+” symbol indicates that PCR amplification was successful, a “**—**” symbol indicates a negative PCR result. nq = DNA below detection limit using the High Sensitivity (HS) double strand (ds) DNA Qubit assays.

| **Site** | **Sample ID** | **DDepth (mbsf)** | **Interpolated in-situ temperature (^o^C)** | **ng DNA per gr of sediment** | **Sediment extracted (gr)** | **Archaeal 16S rRNA primers** | ***mcrA* ANME-1 specific primers** | ***mcrA* general primers (mcrIRD)** |
| --- | --- | --- | --- | --- | --- | --- | --- | --- |
| U1545B | 1545B_1H2 | 1.7 | 5.3 | 300.0 | 0.5 | + | — | — |
|  | 1545B_4H3 | 25.8 | 10.7 | 185.0 | 0.5 | + | + | — |
|  | 1545B_6H2 | 43.6 | 14.7 | 15.8 | 9.5 | + | — | — |
|  | 1545B_11H3 | 92.4 | 25.6 | 3.125 | 8 | + | — | — |
|  | 1545B_13H3 | 111.2 | 29.9 | 3.66 | 5 | + | — | — |
|  | 1545B_15H3 | 130.7 | 34.3 | 3.06 | 5 | + | — | — |
|  | 1545B_20F4 | 160.1 | 40.9 | 2.71 | 5 | + | — | — |
|  | 1545B_25F2 | 177.4 | 44.8 | 1.75 | 5 | + | — | — |
| U1546B | 1546B_1H2 | 0.8 | 2.8 | 900.0 | 0.5 | + | — | — |
|  | 1546B_3H2 | 16.1 | 6.2 | 100 | 0.5 | + | — | — |
|  | 1546B_5H2 | 35.1 | 10.4 | 64.5 | 0.5 | + | — | — |
|  | 1546B_7H2 | 54.0 | 14.6 | 36.2 | 0.5 | + | — | — |
|  | 1546B_9H2 | 73.1 | 18.8 | 16.7 | 0.5 | + | — | — |
|  | 1546B_12H2 | 102.1 | 25.2 | 2.29 | 5 | + | — | — |
|  | 1546B_15H3 | 131.9 | 31.8 | 1.69 | 5 | + | — | — |
|  | 1546B_20H3 | 168.8 | 40.0 | 1.26 | 5 | + | — | — |
| U1547B | 1547B_1H2 | 2.1 | 14.2 | 300 | 0.5 | + | — | — |
|  | 1547B_2H2 | 8.7 | 17.5 | 250 | 0.5 | + | — | + |
|  | 1547B_3H2 | 17.7 | 22.0 | 42.86 | 3.5 | + | — | — |
|  | 1547B_5H2 | 36.9 | 31.9 | 20.0 | 0.5 | + | — | — |
|  | 1547B_7H2 | 55.9 | 41.6 | 2.0 | 5 | + | — | — |
|  | 1547B_8H2 | 65.7 | 46.6 | 2.0 | 5 | + | + | — |
|  | 1547B_9H2 | 74.3 | 51.0 | 0.91 | 5 | + | + | — |
|  | 1547B_12F2 | 94.3 | 60.7 | nq | 5 | + | — | — |
|  | 1547B_25F2 | 132.1 | 80.6 | nq | 5 | + | — | — |
| U1548B | 1548B_1H2 | 2.1 | 8.2 | 700 | 0.5 | + | + | — |
|  | 1548B_2H3 | 8.9 | 13.7 | 135 | 0.5 | + | + | — |
|  | 1548B_3H4 | 20.4 | 22.9 | 33.9 | 0.5 | + | + | + |
|  | 1548B_4H7 | 33.5 | 33.5 | 10 | 0.5 | + | — | — |
|  | 1548B_5H5 | 39.6 | 38.3 | 10 | 0.5 | + | — | — |
|  | 1548B_6H2 | 46.2 | 43.6 | 5.9 | 0.5 | + | — | — |
|  | 1548B_8H2 | 69.5 | 62.4 | 1 | 5 | + | — | — |
|  | 1548B_9H3 | 76.5 | 68.0 | 2.5 | 5 | — | + | — |
| U1549B | 1549B_1H2 | 1.6 | 3.5 | 1040 | 0.5 | + | — | — |
|  | 1549B_2H2 | 7.0 | 4.6 | 364 | 0.5 | + | — | — |
|  | 1549B_3H2 | 16.5 | 6.4 | 161 | 0.5 | + | — | — |
|  | 1549B_6H3 | 45.6 | 12.1 | 63.5 | 0.5 | + | + | — |
|  | 1549B_9H3 | 74.4 | 17.6 | 23.8 | 5 | + | — | — |
|  | 1549B_12H3 | 103.7 | 23.3 | 0.73 | 5 | + | — | — |
|  | 1549B_15H4 | 133.4 | 29.1 | 5 | 5 | + | — | — |
| U1550B | 1550B_1H2 | 2.0 | 3.8 | 1150 | 0.5 | + | + | — |
|  | 1550B_3H2 | 16.9 | 5.8 | 465 | 0.5 | + | + | — |
|  | 1550B_7H2 | 54.8 | 10.9 | 96 | 5 | + | + | + |
|  | 1550B_11H2 | 92.3 | 16.0 | 4.75 | 5 | — | — | — |
|  | 1550B_19X2 | 142.0 | 22.7 | nq | 5 | — | + | — |
| U1551B | 1551B_1H1 | 0.8 | 4.8 | 700 | 0.5 | + | + | — |
|  | 1551B_2H2 | 5.8 | 5.3 | 454 | 0.5 | + | + | — |
|  | 1551B_3H2 | 15.4 | 6.3 | 103 | 0.5 | + | + | — |
|  | 1551B_5H2 | 34.2 | 8.2 | 30 | 0.5 | + | + | — |
| U1552B | 1552B_1H2 | 0.8 | 3.8 | 1100 | 0.5 | + | + | — |
|  | 1552B_3H3 | 19.2 | 8.6 | 605 | 0.5 | + | + | — |
|  | 1552B_3H4 | 20.4 | 8.9 | 129 | 5 | — | — | — |
|  | 1552B_6H2 | 46.9 | 15.8 | 34.4 | 0.5 | + | — | — |

**Supplementary figures**

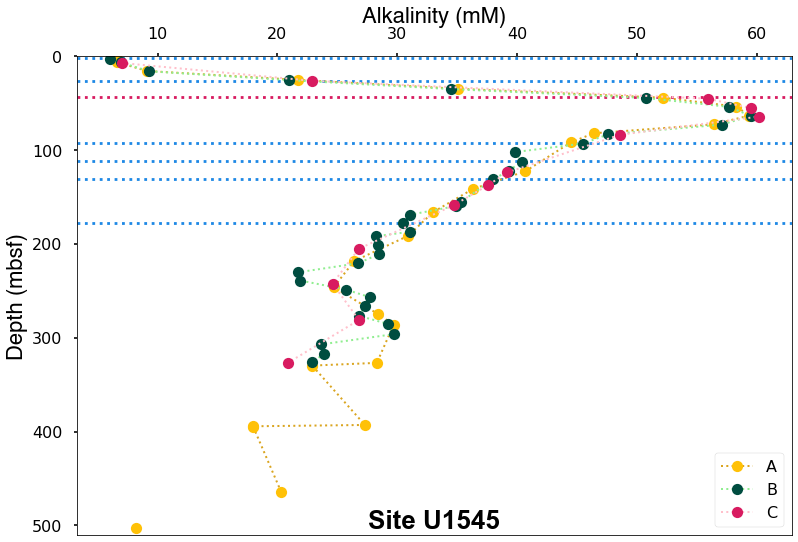

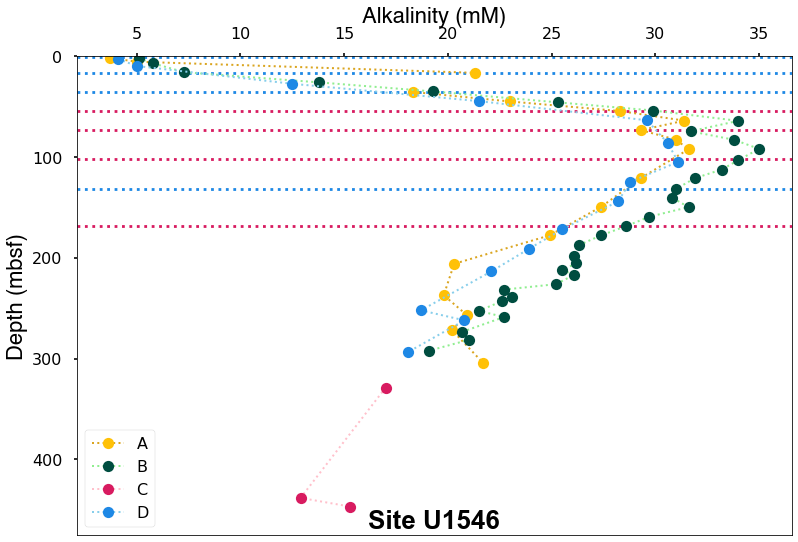

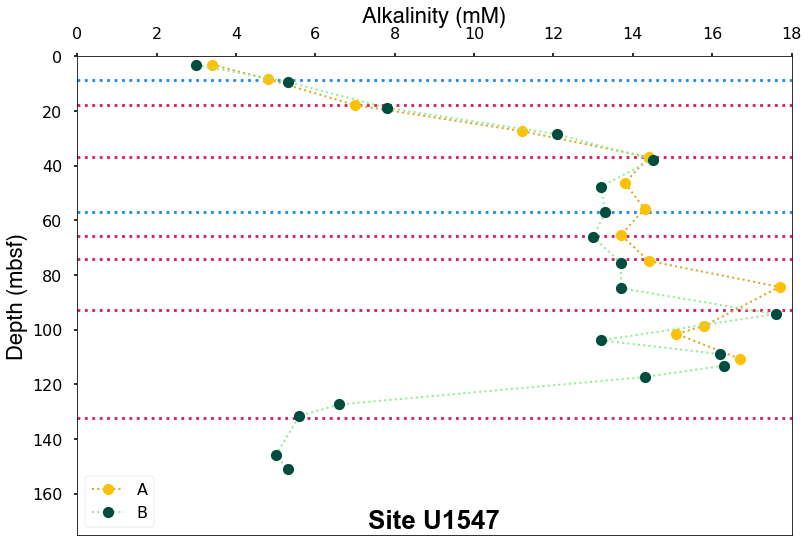

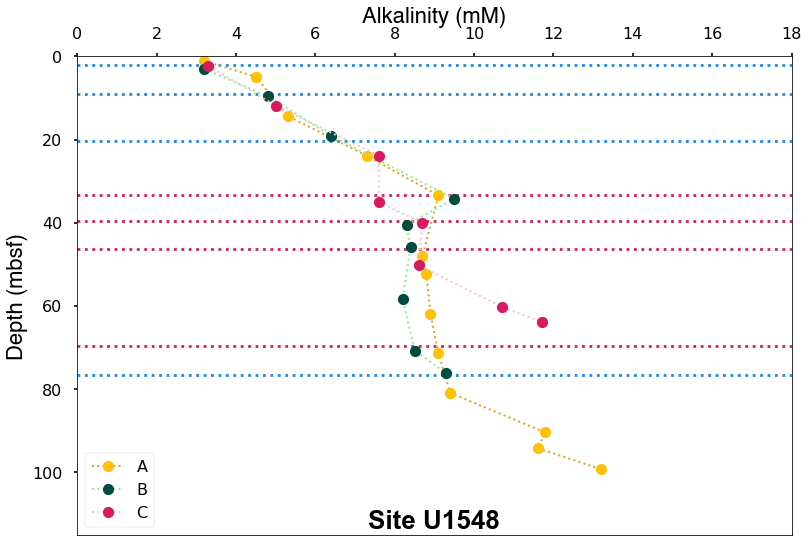

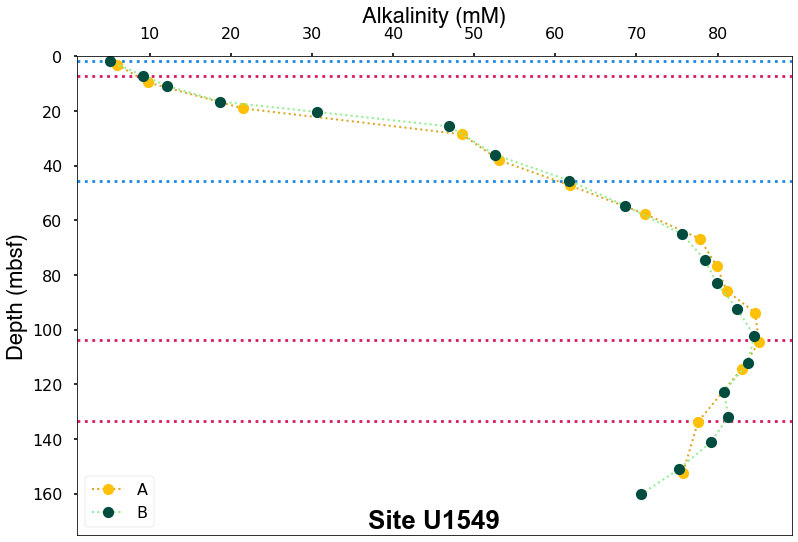

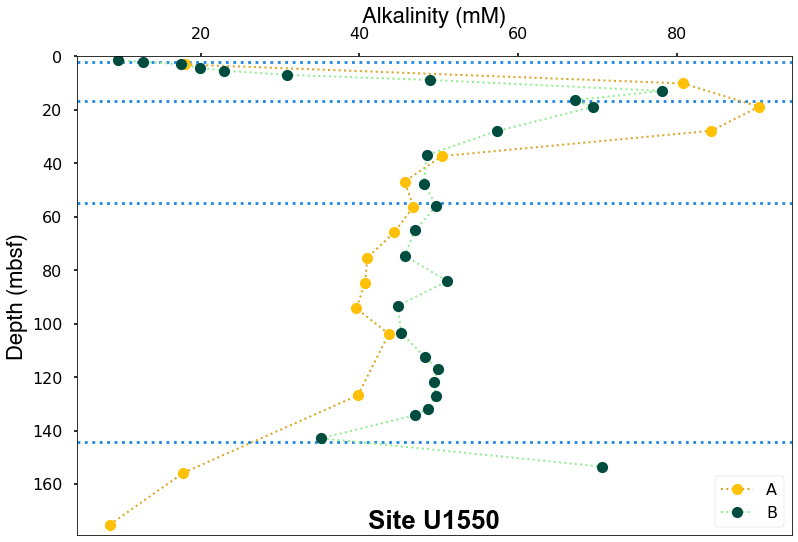

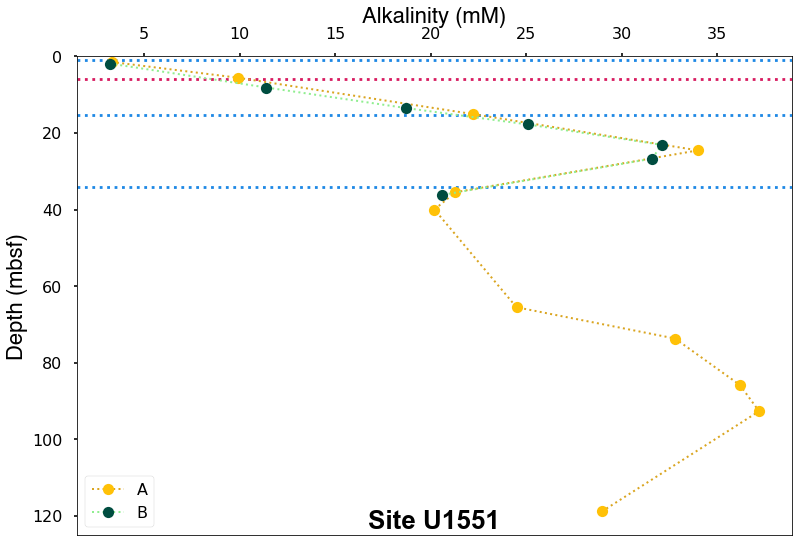

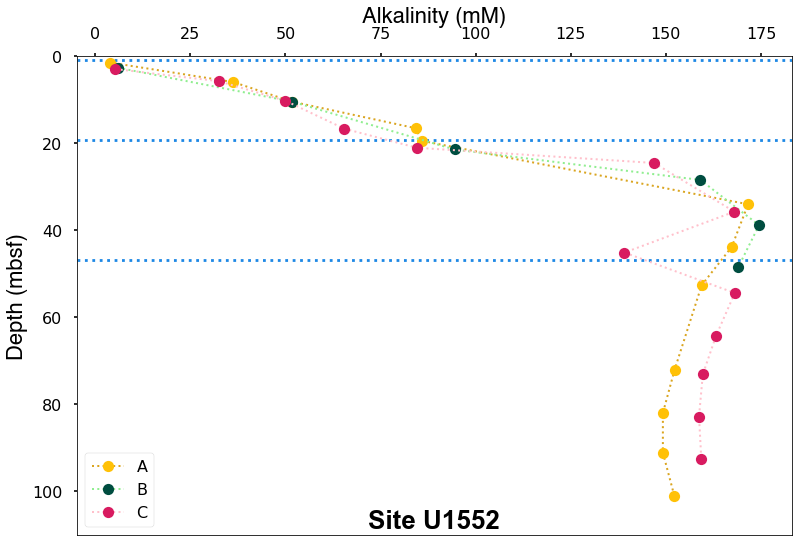

**A**

**B**

**C**

**D**

**E**

**F**

**F)**

**G**

**H**

**Fig. S1**. **Samples and geochemical context.** Sediment horizons that yielded *mcrA* gene amplicons for methanogens and ANME-1 archaea (blue dotted lines) are superimposed on porewater alkalinity gradients from IODP Expedition 385 drilling sites (Teske et al. 2021a-g). Pink lines indicate samples where PCR amplification attempts remained unsuccessful.

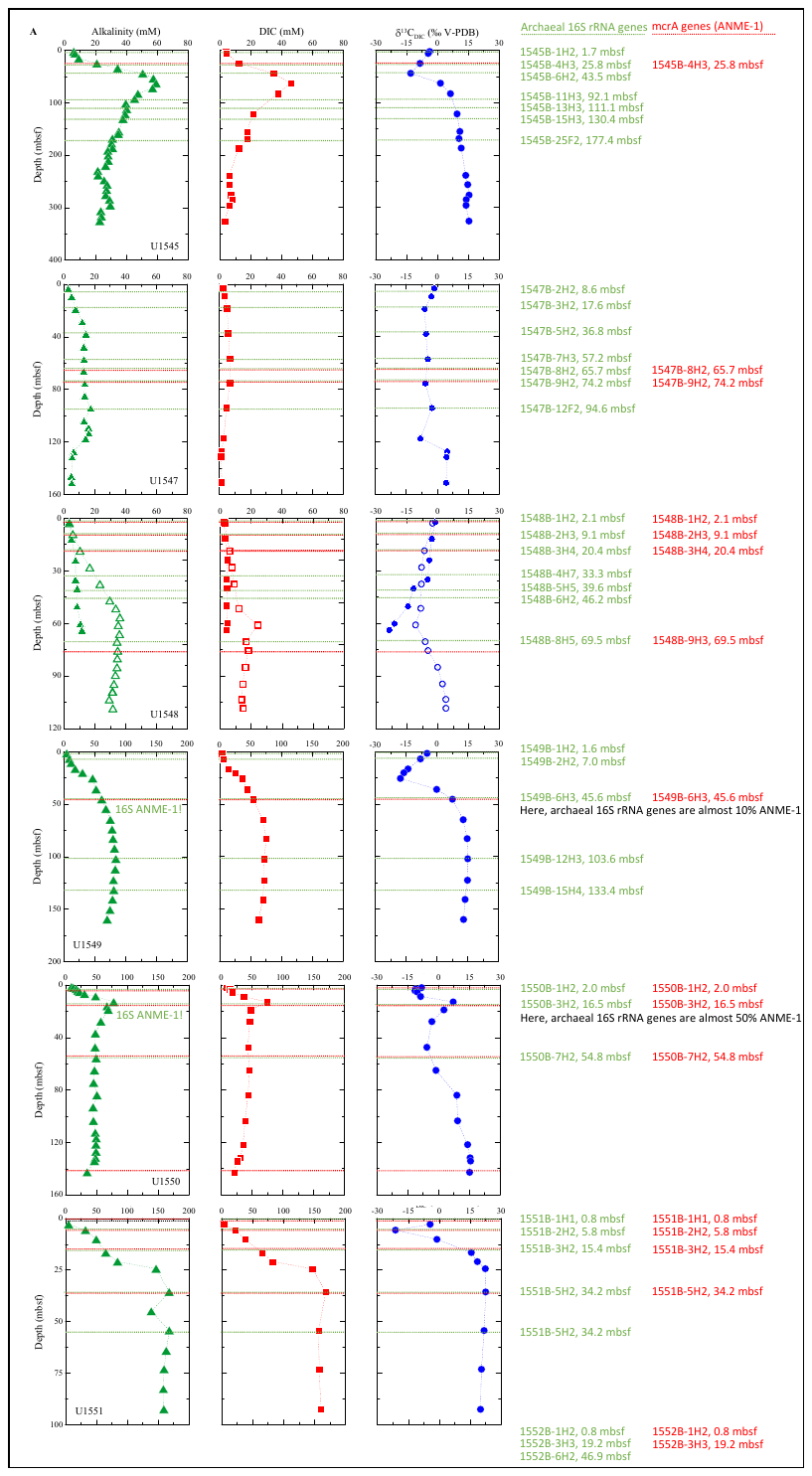

**Figure S2.** **Synopsis of PCR results.** 16S rRNA gene (Mara et al., 2023). and *mcrA* gene detection is superimposed on DIC porewater profiles in IODP 385 sites (Torres and Kim, 2022).

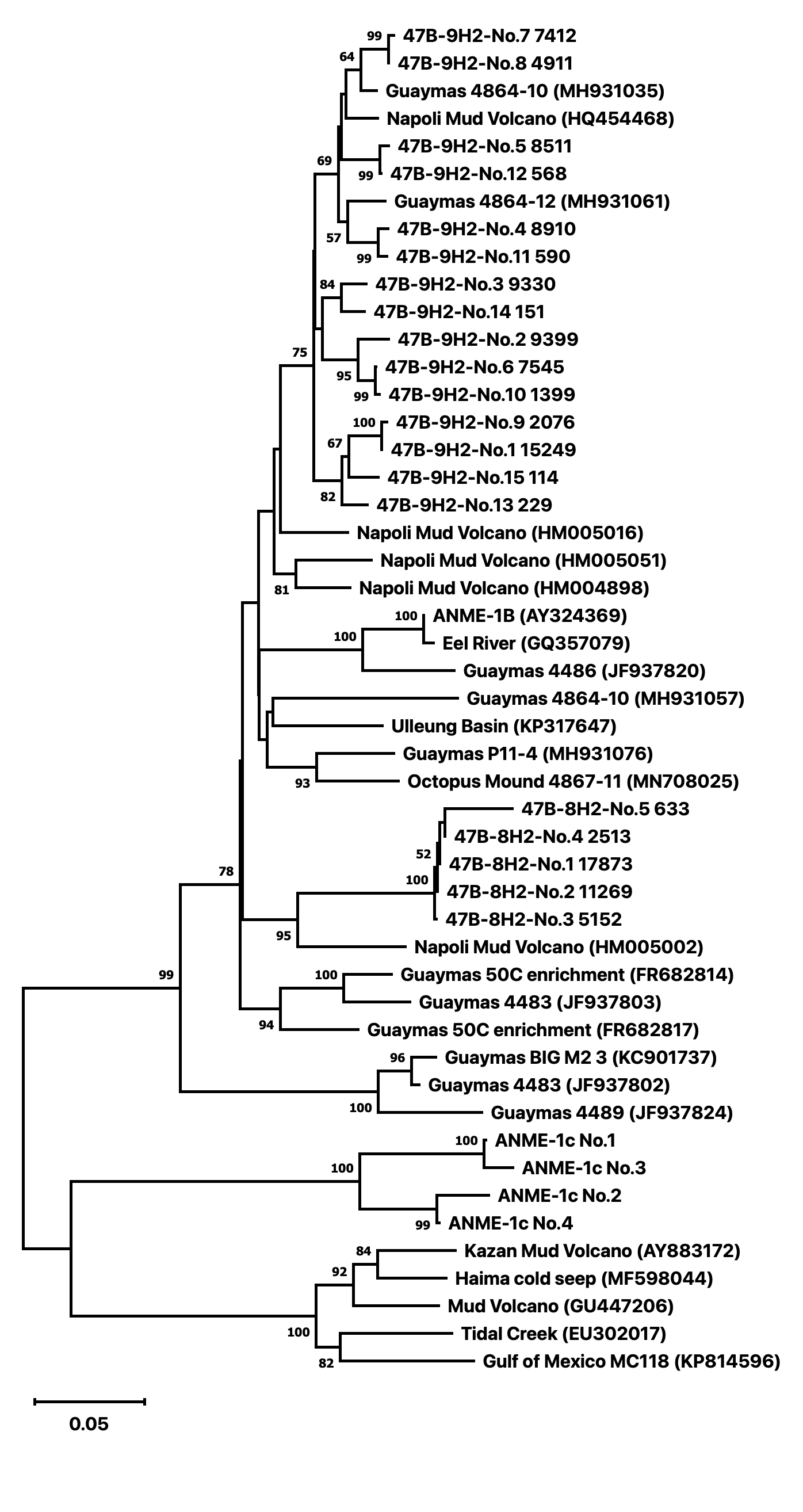

**Cluster III**

**Cluster V**

**Cluster II**

**Cluster XII**

**New Cluster**

**Cluster XI**

**Cluster I**

**Paraphyletic to**

**Cluster I**

**Cluster IX**

**Cluster VI**

**ANME-1c**

**Cluster VIII**

**Cluster XIV**

**Cluster XV**

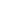

**Cluster III**

**Cluster V**

**Cluster II**

**Cluster XII**

**New Cluster**

**Cluster XI**

**Cluster I**

**Paraphyletic to**

**Cluster I**

**Cluster IX**

**Cluster VI**

**ANME-1c**

**Cluster VIII**

**Cluster XIV**

**Cluster XV**

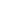

**Figure S3**. Distance (minimum evolution) phylogeny for *mcrA* amplicons from site U1547B (47B), with bootstrap values > 50%. Taxon labels starts with the drilling site (47B), the core number and segment, followed by the ASV number and the number of sequences within each ASV.

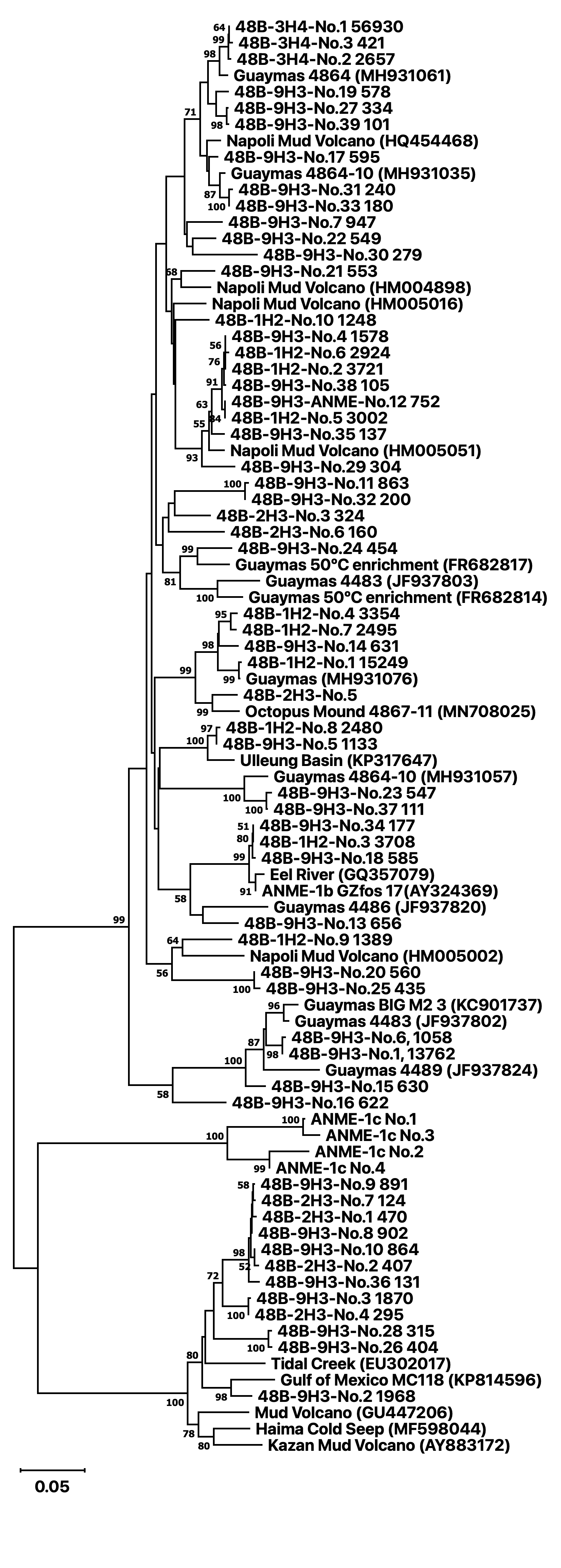

**Cluster XVI**

**Cluster IV**

**Cluster V**

**Cluster VI**

**Cluster XIV**

**XVd**

**XVe**

**XVc**

**XVb**

**XVa**

**Cluster XV**

**ANME-1c**

**IIIb**

**Cluster III**

**IIIe**

**IIId**

**Cluster VIII**

**Cluster II**

**Cluster I**

**Figure S4.** Distance (minimum evolution) phylogeny for *mcrA* amplicons from site U1548B (48B), with bootstrap values > 50%. Taxon labels starts with the drilling site (48B), core number and segment, followed by the ASV number and the number of sequences within each ASV.

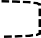

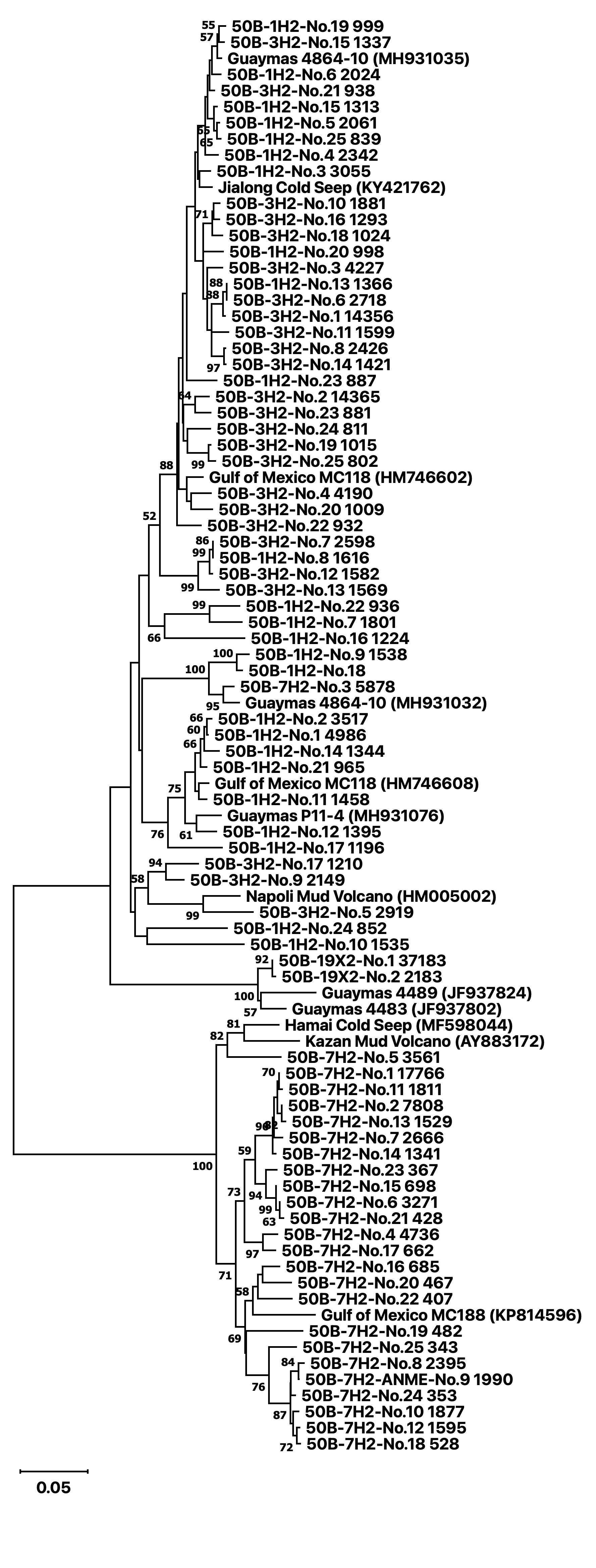

**Cluster XV**

**XVc**

**XVa**

**XVb**

**XVd**

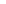

**Cluster XIV**

**Cluster VI**

**Cluster III**

**Cluster VII**

**Cluster XII**

**Cluster X**

**Cluster XI**

**Cluster IX**

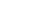

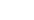

**Figure S5.** Distance (minimum evolution) phylogeny for *mcrA* amplicons from site U1550B (50B), with bootstrap values > 50%. Taxon labels starts with the drilling site (50B), core number and segment, followed by the ASV number and the number of sequences within each ASV.

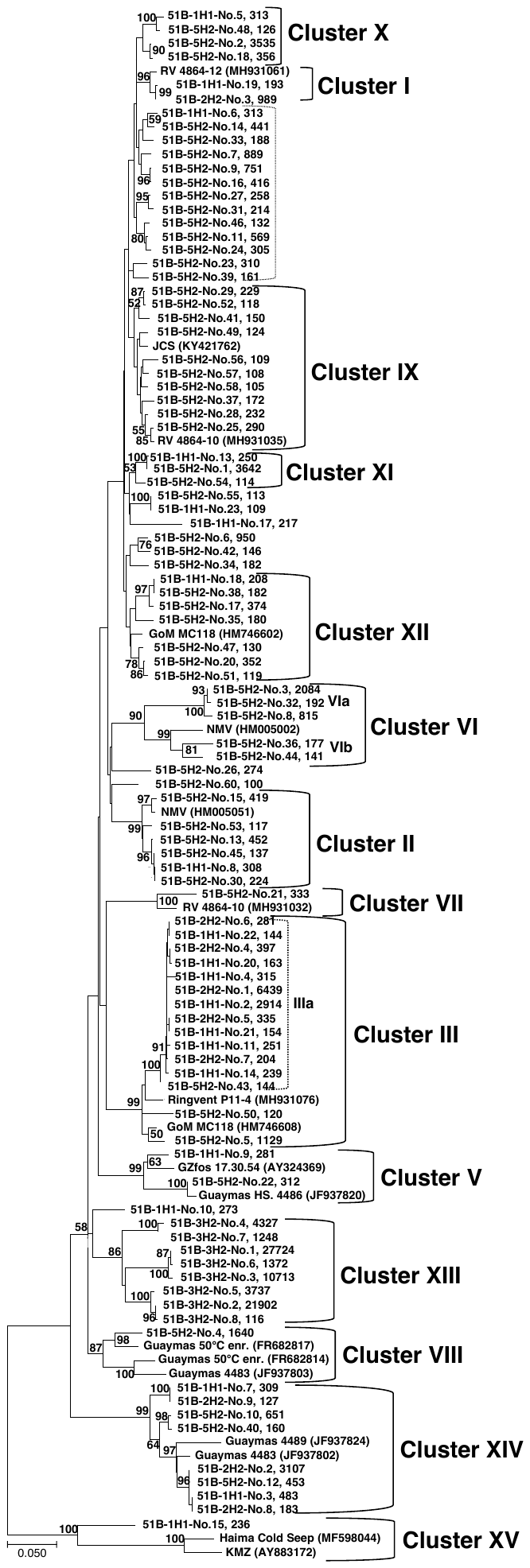

**Figure S6.** Distance (minimum evolution) phylogeny for *mcrA* amplicons from site U1551B (51B), with bootstrap values > 50%. Taxon labels starts with the drilling site (51B), core number and segment, followed by the ASV number and the number of sequences within each ASV.

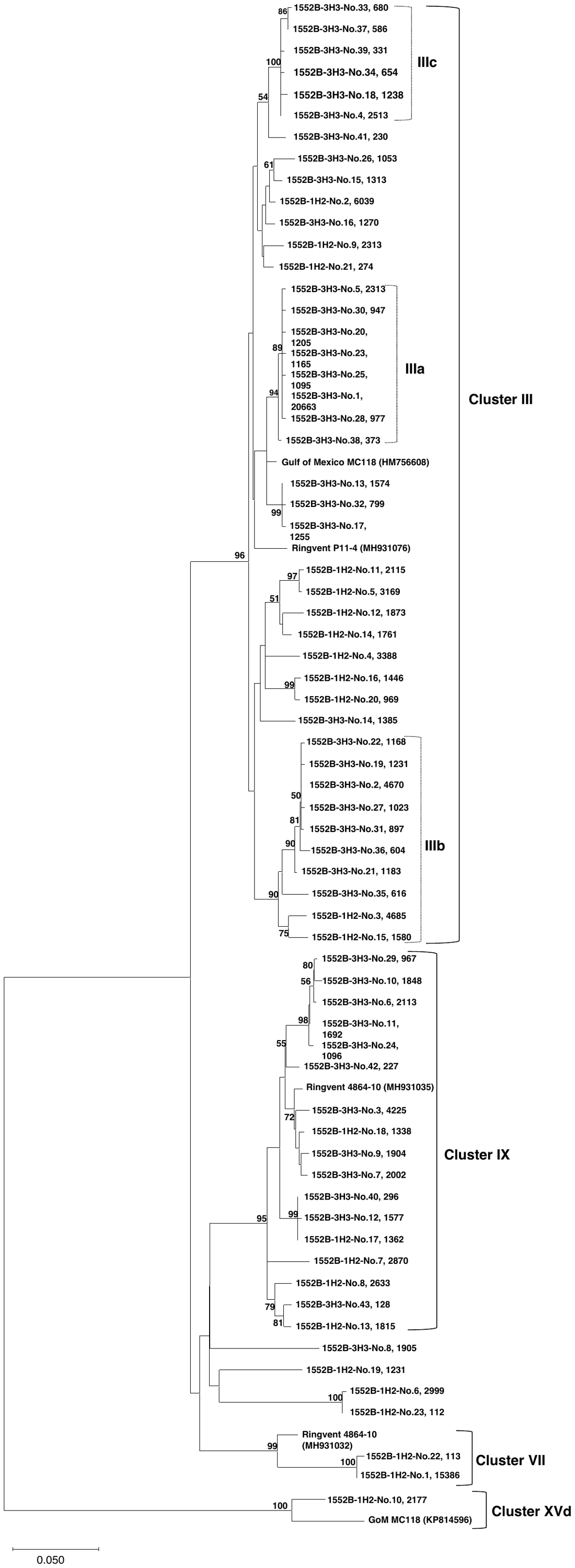

**Figure S7.** Distance (minimum evolution) phylogeny *mcrA* amplicons from site U1552B, with bootstrap values > 50%. Taxon labels starts with the drilling site (1552B), core number and segment), followed by the ASV number and the number of sequences within each ASV.

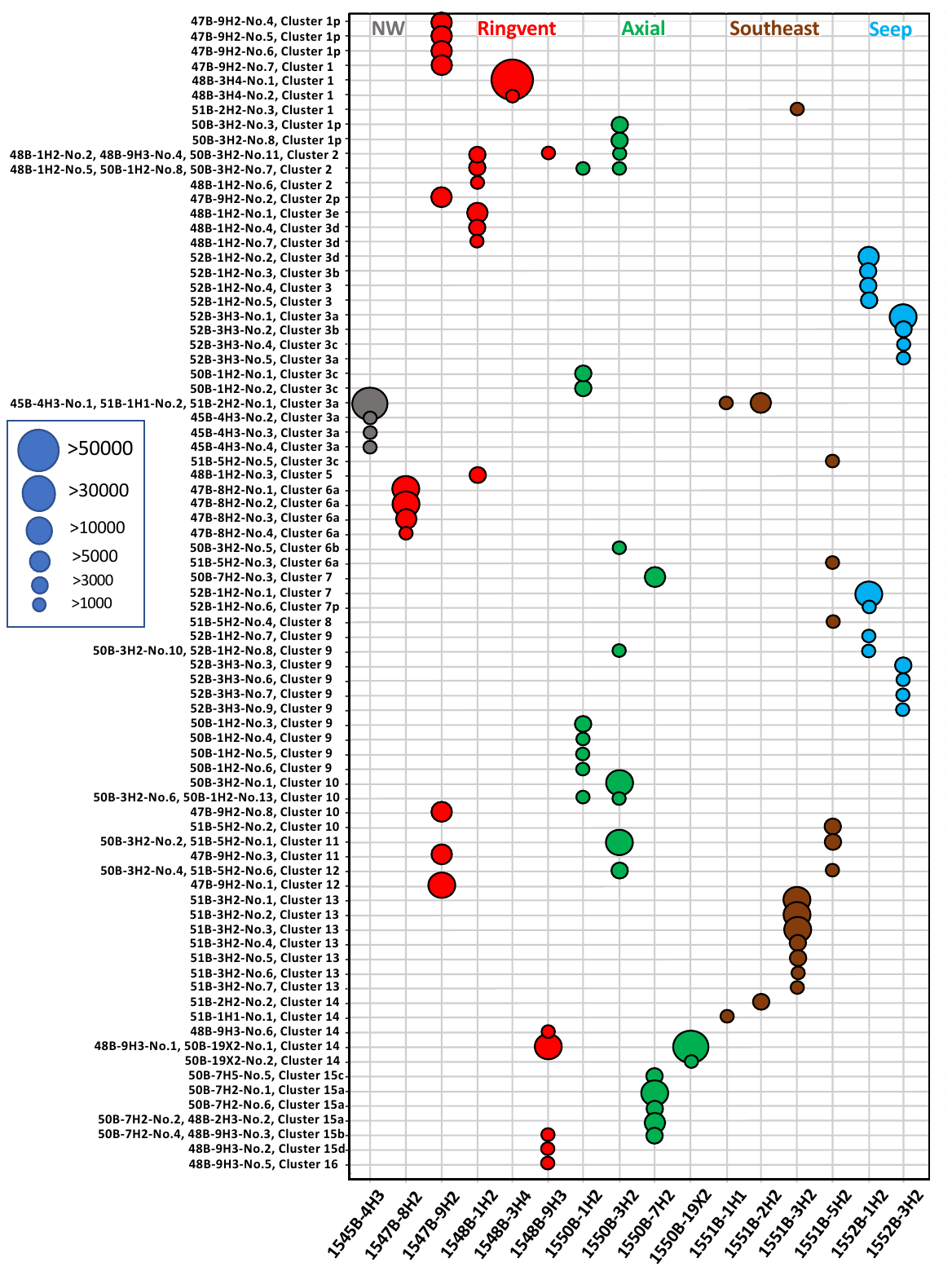
**Figure S8.** Abundance dot plot of *mcrA* gene amplicons (represented by > 1000 sequences) color-coded by site and annotated by sediment sample. Dot size indicates the number of sequences. The *x*-axis lists the samples, and the *y*-axis lists the *mcrA* sequences and their clustering. Suffix “p” indicates the ASV is paraphyletic to an established cluster. These sequences are provisionally included in a cluster but lack strong bootstrap support.
